# The *Tofu* mutation restores female fertility to *Drosophila* with a null *BEAF* mutation

**DOI:** 10.1101/2024.02.13.580197

**Authors:** J. Keller McKowen, Maheshi Dassanayake, Craig M. Hart

## Abstract

Compensatory mutations offer clues to deciphering the role of a particular protein in cellular processes. Here we investigate an unknown compensatory mutation, present in the *BEAF*^*NP6377*^ fly line, that provides sufficient rescue of the defective ovary phenotype caused by null *BEAF* alleles to allow maintenance of fly stocks lacking the chromatin domain insulator proteins Boundary Element-Associated Factors BEAF-32A and BEAF-32B. We call this mutation *Tofu*. We employ both classical genetics and genomic sequencing to attempt to identify the mutation. We find evidence that points to a mutation in a predicted Polycomb response element upstream of the *ribbon* gene, which may lead to aberrant *rib* expression.

## INTRODUCTION

BEAF (Boundary Element-Associated Factor of 32 kDa) is an insulator binding protein of *Drosophila melanogaster* that mainly binds in promoter regions of housekeeping genes (Cuvier et al., 1998; Jiang et al., 2009; Zhao et al., 1995). The *BEAF* gene is located on the second chromosome. It uses alternative promoters to make two isoforms: BEAF-32A and BEAF-32B. Each isoform has a unique N-terminal BED finger DNA binding domain of 81 (32A) or 80 (32B) amino acids (Aravind, 2000). The C-terminal 202 amino acids are common to both isoforms (Hart et al., 1997). This contains a C-terminal BESS domain that mediates BEAF-BEAF interactions, with a leucine zipper immediately adjacent to the BESS domain that strengthens these interactions (Avva and Hart, 2016; Delattre et al., 2002). Previously, our lab generated and characterized a null allele of *BEAF* by eliminating the ATG start codon of both the 32A and 32B specific exons and introducing two tandem stop codons into the shared exon, *BEAF*^*AB-KO*^ (*ABKO*). Flies homozygous for the *ABKO* allele are sickly, display a mild rough eye phenotype, and have defects in oogenesis while males remain fertile. A 32B transgene rescues these phenotypes, while a 32A transgene does not. Ovaries of *ABKO* flies have malformed egg chambers, displaying variable phenotypes that are most apparent at stage eight of oogenesis. As a result, very few mature oocytes are produced. Because of the sharp loss of fecundity, *ABKO* homozygotes cannot be maintained as a stock (Roy et al., 2007).

A separate null allele, *BEAF*^*NP6377*^ (*NP6377*), was generated by the insertion of a GAL4 enhancer trap element containing a mini-*white* marker gene, *P(GawB)*, near the 5’ end of the exon shared by the 32A and 32B isoforms (Hayashi et al., 2002). Thus, in contrast to the *ABKO* allele the *NP6377* allele is marked by *w*. This *NP6377* line was used in a study that attributed the striking phenotypes of neoplastic growth and recessive larval lethality to the loss of BEAF (Gurudatta et al., 2012), which were not observed in our *ABKO* mutant line. To investigate, we obtained the balanced *NP6377* line. We were not able to rescue the phenotypes with two different *BEAF* transgenes that were able to rescue *ABKO* flies (Hart, 2014). The balanced *NP6377* and *ABKO* lines were also crossed in a complementation test, since they were independently generated and so should not have the same second site mutations. The test showed a rescue of the recessive larval lethal and neoplastic growth phenotypes, suggesting they were not a consequence of the loss of BEAF. Surprisingly, *NP6377/ABKO* flies were viable although their ovaries were not completely normal (Fig. 1). The defect in fertility caused by the lack of BEAF was suppressed by a dominant mutation from the balanced *NP6377* line (Hart, 2014). We named this mutation *Tofu* because it allows fly stocks to be maintained without BEAF. By meiotic recombination in the *NP6377/ABKO* line we generated *ABKO/ABKO* (*Tofu KO*) and *NP6377/NP6377* (*Tofu NP)* lines. The recombinant lines lacked BEAF and any recessive lethal mutations, but they retained the dominant enhancer of fertility mutation. Here we describe attempts to locate *Tofu* using two methods: classical genetic mapping via meiotic recombination and high-throughput genomic DNA sequencing.

**Figure 1:**
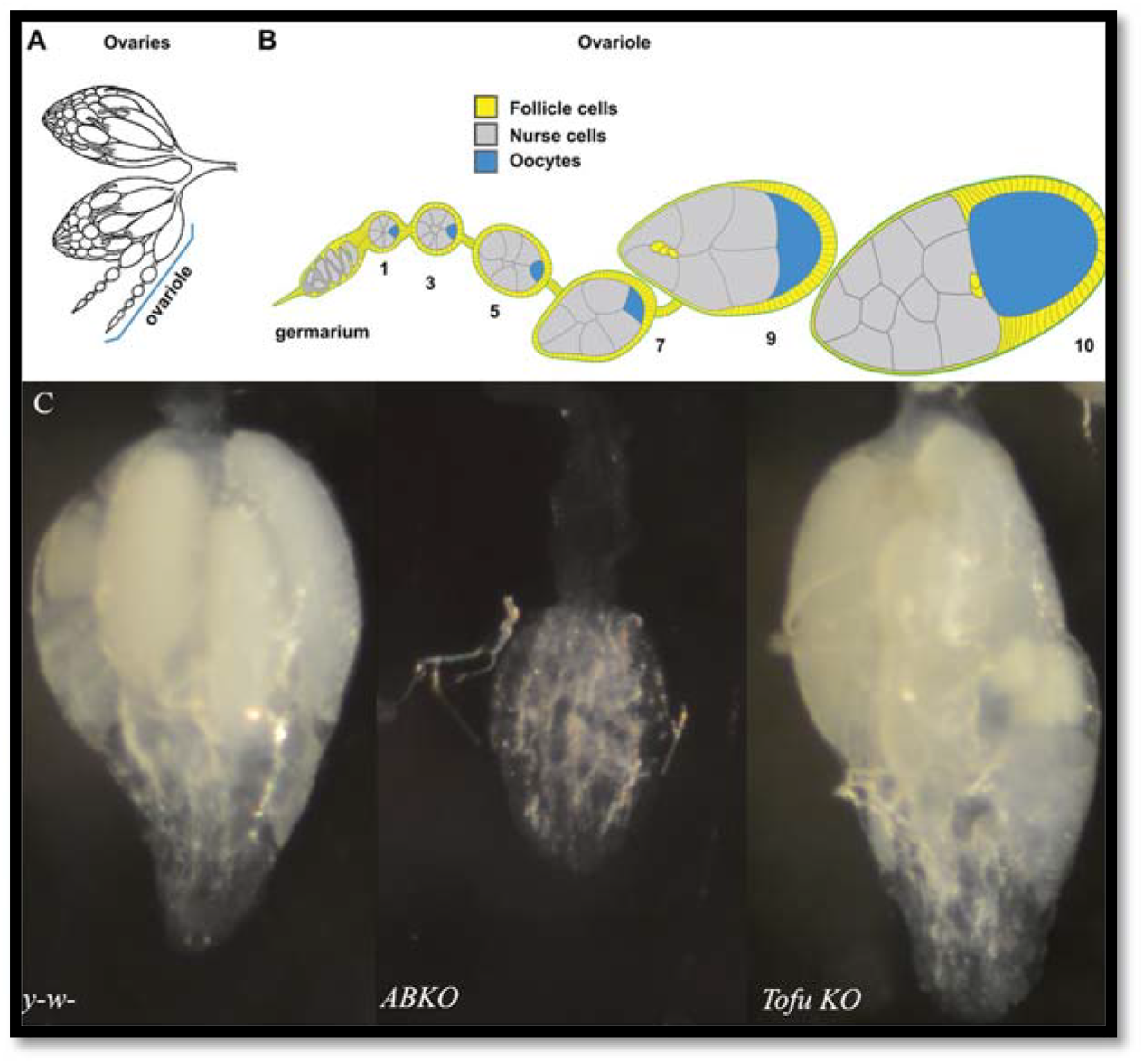
Rescue of the *BEAF*^*AB-KO*^ ovary phenotype by *Tofu*. A: Line drawing of typical fly ovary morphology. Ovaries are composed of several ovarioles which each produce oocytes. B: Oocyte development proceeds from the stem cells at the tip of the germarium to egg chambers, which can be classified as stages by phenotype. Stage 14 is the mature egg (not shown). C: Ovary phenotypes of *y*^*-*^ *w*^*-*^, *ABKO*, and *Tofu KO* flies. *Tofu KO* ovaries have mature eggs, but are atypical. [A & B reproduced with permission from (Andersen and Horne-Badovinac, 2016)]

## MATERIALS AND METHODS

### Fly Stocks

Flies were maintained on standard cornmeal, yeast, and sugar medium with Tegosept. *BEAF*^*NP6377*^*/CyO GFP* fly stocks were kindly provided by Victor Corces (RRID: DGGR 05221). *BEAF*^*AB-KO*^*/ CyO GFP* fly stocks (*CyO GFP* from RRID BDSC 5194) were previously generated by our lab (Roy et al., 2007). *Tofu BEAF* ^*N6377P*^ and *Tofu BEAF*^*AB-KO*^ lines were created by meiotic recombination from a *BEAF*^*AB-KO*^/*BEAF*^*NP6377*^ stock (Hart, 2014). Baylor *P* mapping kit lines (2L: 27E6, 30C1, 33A2, 34B4, 35D2, 36E3, 37B13; 2R: 41D4, 43E11, 47A11, 49E1, 53D4, 55F8, 57E9), *P[w*^*+*^] at 56C6 (RRID: BDSC 18385) and 57B16 (RRID: BDSC 16087), and balancer chromosome lines were obtained from the Bloomington *Drosophila* Stock Center.

### Fly Genotyping

To detect the *ABKO* allele, we designed a genotyping primer set consisting of three primers. The BEAF-wt-Bam-5 has 5’ homology to the *Bam*HI site in the WT *BEAF* allele. The BEAF-mut-5 primer has 5’ homology with the stop codon mutation introduced into the *Bam*HI site of the *ABKO* allele (Roy et al., 2007). The 3’ BEAF-stop+89-3 primer was used with both of these primers to generate a roughly 500 bp product from the corresponding *BEAF* allele.

Genotyping was done using PHIRE tissue direct master mix (Thermo-Fisher).

BEAF-wt-Bam-5: GACATCATATACAGCGAGGATCC

BEAF-mut-Bam-5: AAGGACATCATATACAGCGAGTAATG

BEAF-stop+89-3: TTACGACACGCTGATTTGCC

### Genomic DNA preparation

Fifty wandering third instar larvae from y^-^ w^-^, *ABKO/CyO-GFP, NP6677/CyO-GFP, Tofu KO, and Tofu NP* lines were used for DNA isolation. Larvae homozygous for *BEAF* knockout alleles were selected from the *ABKO/CyO-GFP*, and *NP6677/CyO-GFP* lines by lack of GFP expression. Larvae were homogenized in 500 μL of Buffer A (10 mM Tris-Cl pH 7.5, 60 mM NaCl, 10 mM EDTA, 150 μM spermine, 150 μM spermidine, 200 μg/mL Proteinase K). Then 500 μL of Buffer B (200 mM Tris-Cl (pH 7.5), 30 mM EDTA, 2% SDS, 200 μg/mL Proteinase K) was added, and the samples were incubated for one hour at 37° C. Samples were purified by phenol extraction, then phenol-chloroform-isoamyl alcohol (25:24:1) extraction, and then chloroform extraction. Samples were ethanol precipitated and dissolved in 10 mM Tris-Cl pH 7.5.

### Illumina Sequencing

Genomic DNA was submitted to the Roy J. Carver Biotechnology Center at the University of Illinois at Urbana. Libraries were prepared from 300 bp to 500 bp fragments for each sample followed by paired end sequencing of 150 bp from each end using an Illumina HiSeq2500 instrument.

### Data analysis

Raw reads were obtained as paired fastq files (deposited at NCBI BioProject accession number PRJNA1069327 ). Read quality was analyzed using FASTQC (Andrews, 2010). Reads were aligned to the dmel r6.09 genome using bowtie2. Samtools was used to assess alignment quality and compress sam alignments into bam format. Variant calling was done using the GATK toolkit (DePristo et al., 2011; McKenna et al., 2010; Van der Auwera et al., 2013). Variant call files (VCF) were compared using bedTools (Quinlan, 2014). Protein coding changes were called with SnpEff (Cingolani et al., 2012). SNP density was visualized as bigwig files using 2000 bp windows. File conversion was done using Unix commands, Kent utilities (Kent et al., 2010), bedTools, and deepTools (Ramirez et al., 2016). Data was displayed using IGV (Robinson et al., 2011).

## RESULTS

### *Tofu* is on Chromosome 2

Discovery of the *Tofu* allele indicated it was on the same chromosome as *BEAF*, chromosome 2 (Hart, 2014). To check this, we performed a chromosome segregation analysis. In the first strategy (Fig. 2), *Tofu KO* flies were mated with a *CyO/Wg*^*Sp-1*^*;TM3/Scm*^*ET50*^ double balancer line. We assayed 200 females from the second cross and found a fertility rate of 53%. This indicates *Tofu* is not on chromosome 3, but it does not clearly support association with chromosome 2. *Tofu* could be on chromosome 2 if it was at least partially heterozygous in the *Tofu KO* stock and/or some of the single pair mating vials were scored as infertile because the vials were not healthy. The incomplete rescue of ovary function (Fig. 1) and other possible unknown fitness issues could also contribute. *Tofu* could also be on chromosome X or 4.

**Figure 2:**
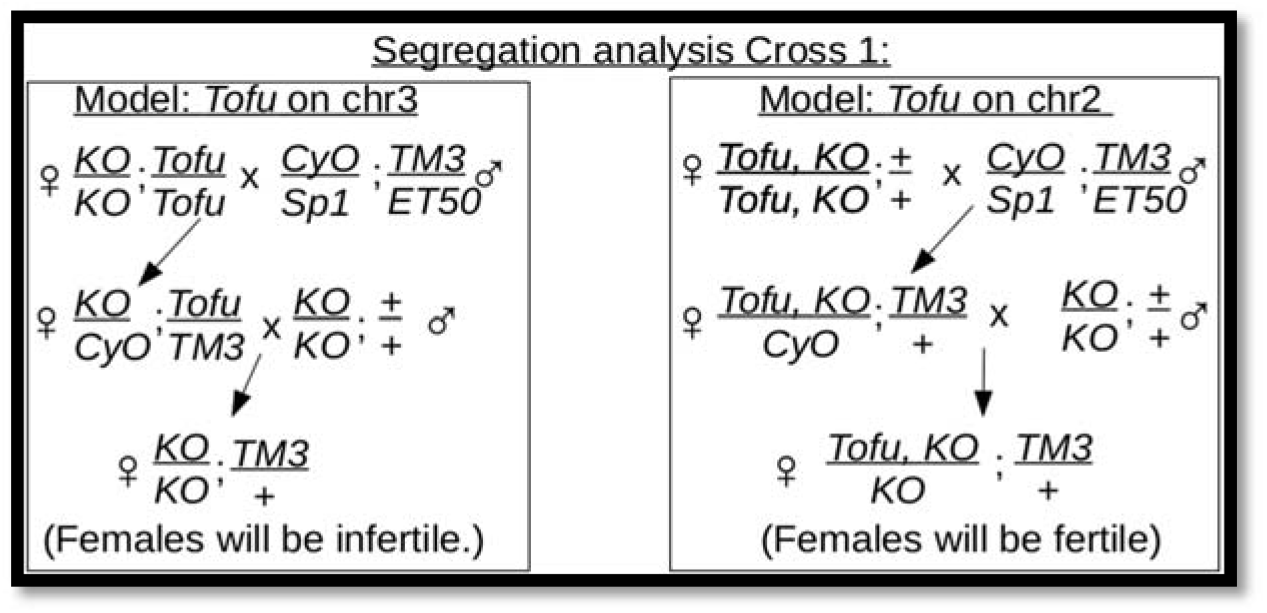
Diagram of a segregation cross for chromosomes 2 and 3. The mating strategy takes advantage of the fertility of *ABKO* males, and is divided into two models of expected results. A female fertility rate of 53% was found, rather than the expected 0% or 100%.

In the second strategy (Fig. 3), males of the *Tofu KO* line were mated to *ABKO/CyO* virgin females. We assayed 200 female progeny from the second cross and found a fertility rate of 52%. This indicates *Tofu* is not on chromosome X and suggests it is associated with chromosome 2. This strategy does not exclude the possibility that *Tofu* is on chromosome 4, although given the small size and low number of genes this is unlikely. Also, single *Tofu KO*/*CyO* and *Tofu NP*/*CyO* flies could be crossed to *CyO*/*Sp1* for several generations and retain the ability to make *Tofu KO* and *Tofu NP* homozygous lines. This suggests that *Tofu* is on the same chromosome as the *BEAF* alleles.

**Figure 3:**
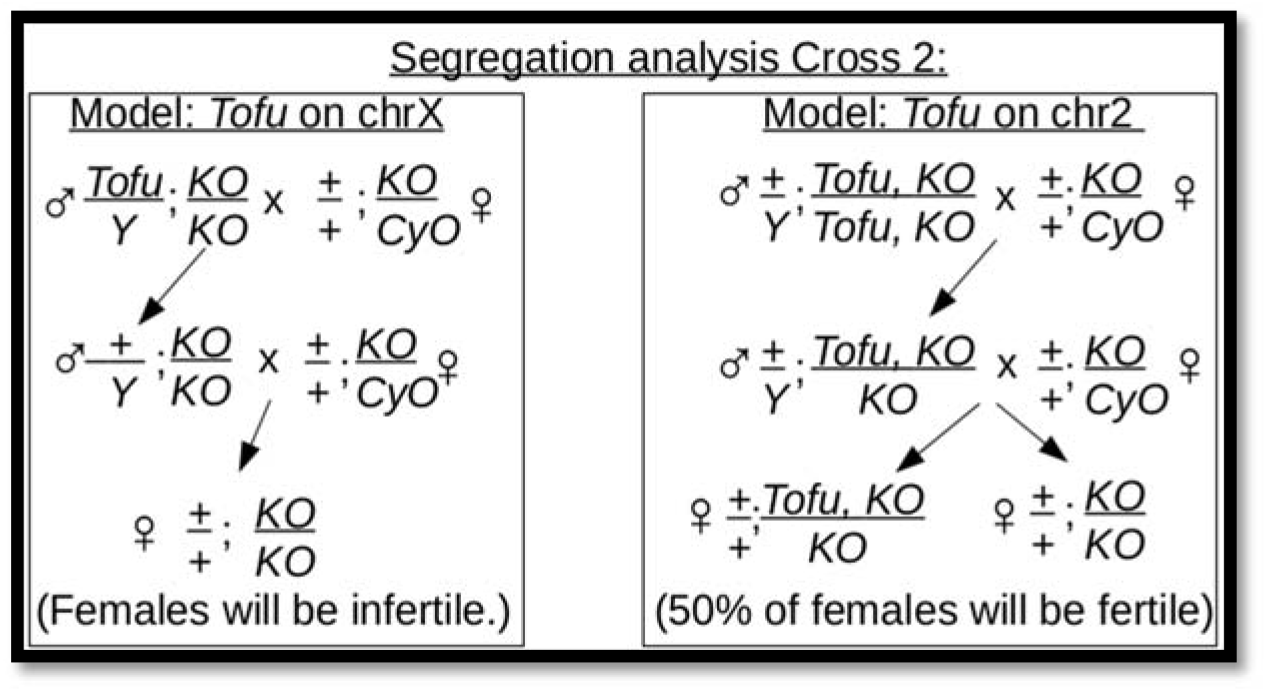
Diagram of a segregation cross for chromosomes X and 2. The mating strategy is divided into two models of expected results. A female fertility rate of 52% was found.

### Mapping by Meiotic Recombination

To narrow down a region that contains *Tofu*, we employed meiotic recombination in females to map genetic linkage. Fly lines with chromosome 2 *w*^*+*^ P element insertions at known cytological locations from the Baylor P mapping kit were used (Zhai et al., 2003). The *ABKO* allele was recombined onto the *P[w*^*+*^] chromosome and confirmed by PCR (Fig. 4A). This chromosome was then used in the strategy shown in Figure 4B to get the recombination frequency of *Tofu* off of the *Tofu ABKO* chromosome and onto the *P[w+] ABKO* chromosome, using fertility of individual females as the phenotype. If *Tofu* recombined onto the *P[w*^*+*^] chromosome, females with red eyes should be fertile and with white eyes should be infertile. For each *P[w*^*+*^] line we scored 100 single-pair matings for females with red eyes and 100 for females with white eyes. The anticipated result was that the percentage of fertile females with red eyes would equal the percentage of infertile females with white eyes, and the percentage would be much less than 50% if *P[w*^*+*^] and *Tofu* were cytologically close. These single-pair matings proved to be unreliable (Fig. 4C), possibly because of unknown genetic interactions with genes from the *P[w*^*+*^] mapping lines or unhealthy environmental conditions in some single-pair mating vials.

**Figure 4:**
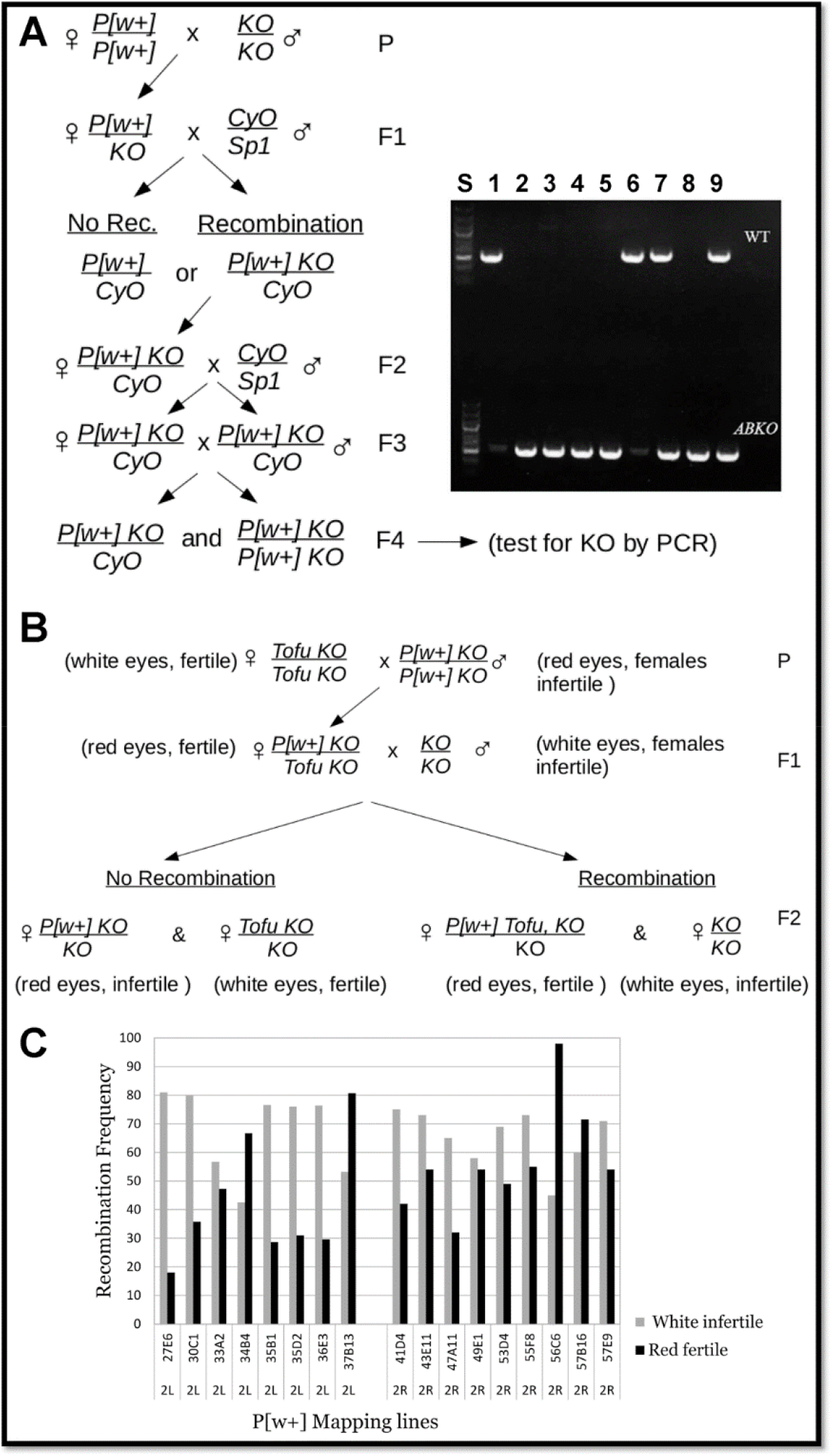
Meiotic recombination mapping strategy. (A) Strategy for recombining *P[w*^*+*^] onto the *ABKO* chromosome, and PCR detection of recombination for 27E6 (535 bp PCR product for *wt* and *ABKO*). Agarose gel lanes: S: standard (lower bright band is 500 bp); 1: *wt* fly; 2-8: recombination candidate flies (homozygous males except lane 7 is a female with *CyO*); lane 9: *ABKO*/*CyO* fly. PCR used primers for the *wt* allele (upper tier) or *ABKO* allele (lower tier). (B) Mapping strategy. (C) Flies from the mapping strategy were segregated based on eye color and individual females were scored for fertility (White: n=100; Red: n=100).

We tested our ability to do meiotic mapping using two strategies. This also allowed us to determine recombination frequencies of cytological locations relative to the *BEAF* gene near the center of chromosome arm 2R at 51C2. First we used the recessive marker mutation *speck* (*sp*) which is in the telomere-proximal 60B12-60C1 region of 2R (∼9.3 Mb from *BEAF*). This mutation is present in our *ABKO* and *Tofu KO* lines, but not in the *NP* lines. Using the strategies shown in Figure 5A and B, we found that *sp* recombined onto the chromosome with *NP* with a frequency of 49% (n=305) and off again with a frequency of 56% (n=111). Second, we used P mapping lines with *ABKO* recombined onto the two chromosomes with *P[w*^*+*^] inserted nearest to *BEAF*, 49E1 and 53D4 (∼1.8 Mb and 1.5 Mb from *BEAF*, respectively). Using the strategy shown in Figure 5C and PCR to determine if the *ABKO* allele was present, we found that *ABKO* separated from the 49E1 *P[w*^*+*^] with a frequency of 7.6% (n=79) and from 53D4 *P[w*^*+*^] with a frequency of 11.3% (n=71). The results demonstrated our ability to identify linked regions, confirming that the failure of our fertility assay was due to problems accurately scoring fertility.

**Figure 5:**
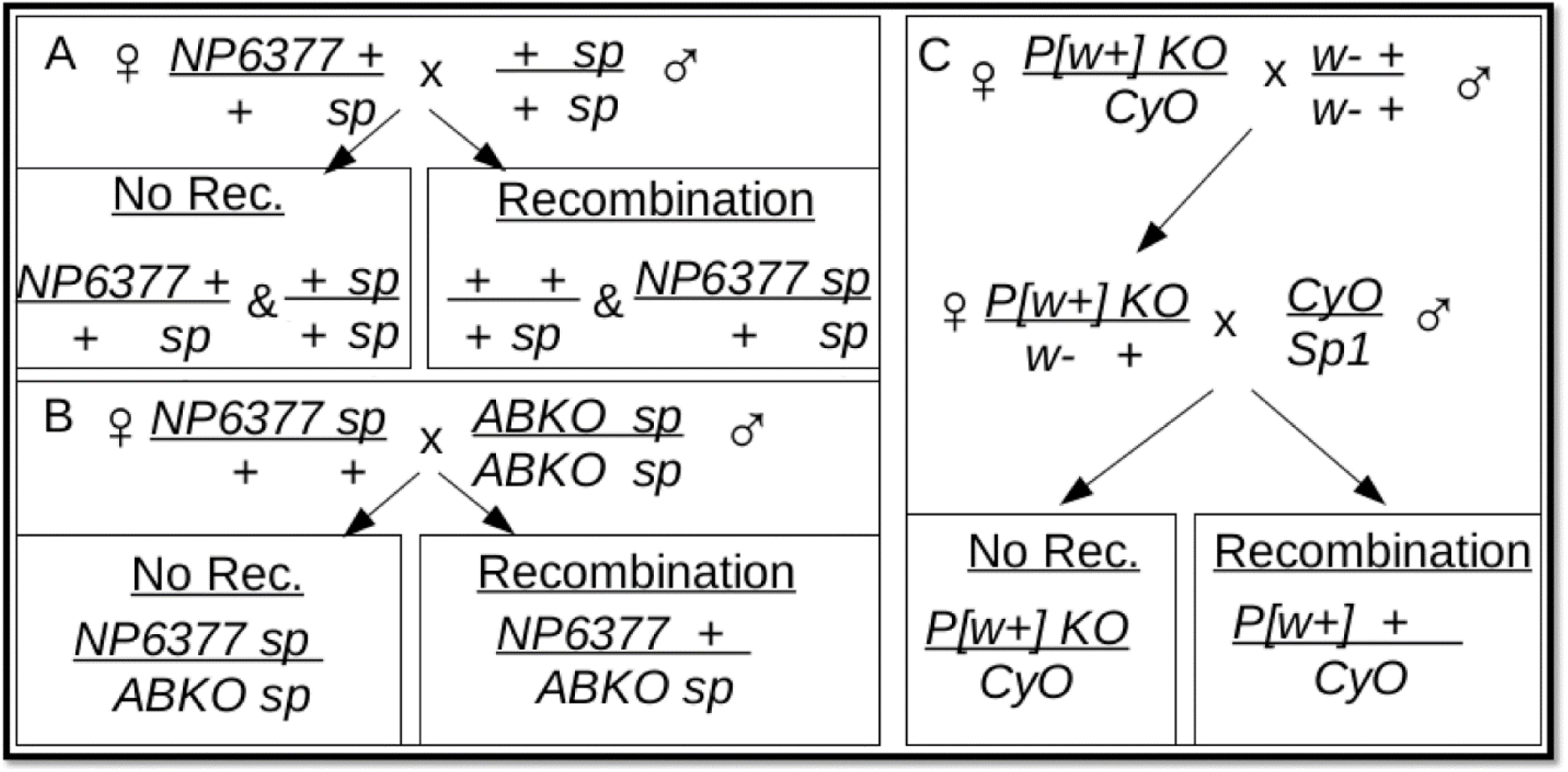
Meiotic recombination mapping strategy for *BEAF* alleles at 51C2 relative to *speck* at 60B12-60C1 and *P[w*^*+*^] at 49E1 and 53D4. (A) Strategy for detecting recombination of *speck* (*sp*) onto the *NP6377* chromosome. A recombination rate of 49% was found (n=305). The *w*^*+*^ transgene in the *NP6377 BEAF* allele was used for identification. (B) Strategy for detecting recombination of *sp* off of the *NP6377* chromosome. A recombination rate of 56% was found (n=111). (C) Strategy for detecting recombination of *P[w*^*+*^] off of the *ABKO* chromosome. Presence or absence of *ABKO* was detected by PCR of flies with red eyes, as in Fig. 4A. A recombination rate of 7.6% was found for 49E1 (n=79) and 11.3% for 53D4 (n=71). Flies in all crosses had a *w*^*-*^ chromosome X.

### Genomic Sequencing

In another approach to identify *Tofu* candidates, we used genomic sequencing to compare mutations among fly lines. Genomic DNA was isolated from homozygous larvae of five lines: *y*^*-*^ *w*^*-*^, *ABKO, NP6377*, and the two lines generated from recombination: *Tofu NP* and *Tofu KO*. The *Tofu* mutation is in *NP6377, Tofu NP*, and *Tofu KO*, but not *ABKO* and *y*^*-*^ *w*^*-*^. Illumina sequencing was used to generate paired end reads of 150 bp per end from 300-500 bp DNA fragments (Bentley et al., 2008). Sequence reads were aligned to the dm6 genome release for *D. melanogaster* using bowtie2 after filtering for quality and trimming adapters, resulting in a genome coverage of over 40 reads per base (Table 1). Paired end sequencing improves allignment accuracy, and an excessive distance between read pairs can indicate an insertion mutation relative to the reference genome (Langmead and Salzberg, 2012; Li et al., 2009).

**Table 1:** Genomic sequencing statistics.

| Sample | Number of Reads Per End | Filtered, Clipped, Paired and Mapped Reads | Mean Quality (PHRED) | Mean Fragment (bp) | Fold Genome Coverage |
| --- | --- | --- | --- | --- | --- |
| **ABKO** | 32,067,905 | 19,177,162 | 33 | 347 | 48 |
| **Tofu KO** | 29,721,768 | 16,853,622 | 36 | 363 | 42 |
| **NP6377** | 31,616,665 | 19,035,482 | 36 | 361 | 47 |
| **Tofu NP** | 28,497,357 | 16,010,932 | 36 | 372 | 40 |
| **y- w-** | 28,514,732 | 16,028,674 | 36 | 355 | 40 |

The aligned genomes were used to call variants relative to the dm6 reference genome by using a suite of programs in the genome analysis toolkit, GATK (DePristo et al., 2011; McKenna et al., 2010; Van der Auwera et al., 2013). The resulting variant call files (VCFs) for each fly line detail the location and sequence change from the reference genome for single nucleotide polymorphisms (SNPs) and extra bases or missing bases (insertions or deletions, lumped together as INDELs) (Supplementary files S1-S5). For simplicity of discussion, we will use SNPs to refer to both SNPs and INDELs. The VCFs for each of the 5 lines had over 700,000 SNPs relative to the reference genome (Table 2). After cross-comparing, we found 61,693 SNPs that fit the pattern for *Tofu* among the lines sequenced (Supplementary file S6). Of these, 1,203 SNPs were localized to chromosome 2, a small number compared to other chromosomes. We reasoned this was due to recombination-mediated shuffling and selection between the original *ABKO* and *NP6377* chromosomes to get *Tofu* onto the *ABKO* chromosome while removing recessive lethal mutations from the *NP6377* chromosome.

**Table 2.** Chromosome arm sizes and SNP counts per chromosome in sequenced lines.

|  | chr2L | chr2R | chr3L | chr3R | chrX | chr4 | Total |
| --- | --- | --- | --- | --- | --- | --- | --- |
| **size (Mb)** | 23.51 | 25.29 | 28.11 | 32.08 | 23.54 | 1.35 | 133.88 |
| **NP6377** | 164,335 | 148,240 | 181,649 | 149,138 | 121,174 | 2,159 | 766,695 |
| **Tofu NP** | 215,087 | 192,164 | 192,112 | 179,544 | 113,948 | 1,872 | 894,727 |
| **Tofu KO** | 168,156 | 145,573 | 191,076 | 179,089 | 106,302 | 1,373 | 791,569 |
| **ABKO** | 166,805 | 145,117 | 229,148 | 186,378 | 119,324 | 1,400 | 848,172 |
| **y- w-** | 167,007 | 148,868 | 200,959 | 150,514 | 104,656 | 1,156 | 773,160 |
| **Tofu Pattern** | 71 | 1,132 | 34,711 | 20,940 | 4,783 | 56 | 61,693 |

Tofu pattern: SNPs present in the *NP6377*, *Tofu NP*, and *Tofu KO* sequences but absent in the *ABKO* and *y- w-* sequences.

We used SnpEff to predict the effect of the chromosome 2 SNPs on protein synthesis and found 35 of the SNPs would cause missense, frameshift, or splicing mutations in 17 protein-coding genes (Supplementary file S7). Only 2 of these genes popped out as likely candidates. One, *18 wheeler* (*18w*), encodes a Toll-like receptor family member and plays a role in various processes including ovarian follicle cell migration (Kleve et al., 2006). However, *18w* has a conservative Ala1357Val mutation near its C-terminus that might not affect function. The other, *hu li tai shao* (*hts*), encodes an adducin homolog that is associated with oocyte fusomes and ring canals and is essential for fertility (Lin et al., 1994; Yue and Spradling, 1992). However, the 3 missense mutations in *hts* (Lys1760Glu, Ile1764Met, Thr1773Asn) affect only 1 of 16 splice variants (*hts-RP*), and this isoform has not been found in ovaries [(Petrella et al., 2007) and FlyBase JBrowse RNA-seq data]. Therefore we focused on the possibility that *Tofu* is not a gain-of-function protein coding mutation.

We re-examined the *Tofu* candidate variants by plotting them using the Integrative Genomics Viewer (IGV) (Robinson et al., 2011). Meiotic recombination between the original *NP6377* and *ABKO* chromosomes generated the *Tofu NP* and *Tofu KO* fly lines by removing one or more recessive lethal mutations from the *NP6377* chromosome and transferring *Tofu* to the *ABKO* chromosome. We reasoned that this process would also transfer linked variants near *Tofu*, so the *Tofu* mutation should lie in a variant dense region. While there are several variant dense regions, only one is on chromosome 2 (Fig. 6). This region on 2R contains 954 of the 1,203 candidate SNPs on the second chromosome, including 32 of the 35 SNPs that changed protein coding sequences. We refer to this region as the “SNP island”.

**Figure 6:**
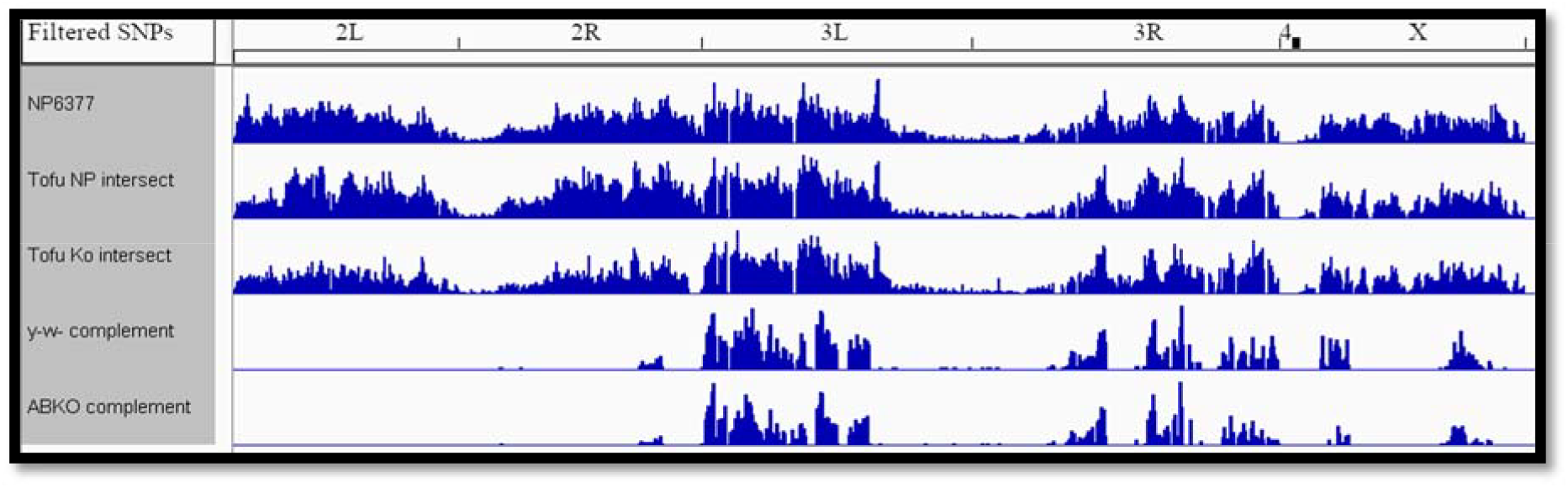
IGV visualization of SNP density. The SNPs are filtered from top (unfiltered) down. The top 3 genomes have *Tofu*, the bottom 2 do not. The bottom row shows candidate SNPs for *Tofu*.

The SNP island corresponds to cytological regions 55F8-57C3 (dm6 chr2R:18,868,928-21,135,871, nearly 2.3 Mb). The original P-element fly lines we used were outside of this region, so we obtained *P[w+]* insertion lines within the SNP island at 56C6 and 57B16. The recombination rates were higher than expected if *Tofu* is in this region (Fig. 4). Because we suspect that *Tofu* is in this region, these results reinforce our reservations about the reliability of the fertility assay.

Rather than a mutation in protein-coding sequences, *Tofu* could be a cis-regulatory element (CRE) mutation. It is often difficult to predict if mutations are in CREs because they do not alter protein coding sequences, and the limited knowledge of transcription factor binding specificity makes it difficult to predict if a SNP will alter factor binding at a given site. Additionally, a given CRE can alter target gene expression from great distances and CRE-promoter specificity is not well characterized for most genes, making it difficult to identify target genes. An exception would be a mutation in a Polycomb response element (PRE). PREs can facilitate the initiation and spreading of a histone H3K27me3 domain, a mark associated with repression of genes located in the domain (Kuroda et al., 2020). A PRE mutation could result in increased transcription of the gene that rescues fertility (gain of function) or expression in new cell types (function in an atypical context), which would be consistent with the *Tofu* mutation. A previous study predicted 537 PREs based on chromatin analysis (Negre et al., 2011). We found only two predicted PREs with candidate SNPs on the second chromosome, and both are within the SNP island region of 2R (Fig. 7A). One is upstream of the gene *Act57B*, while the other is upstream of the *ribbon* (*rib*) BTB/POZ domain transcription factor gene (Fig. 7B).

**Figure 7:**
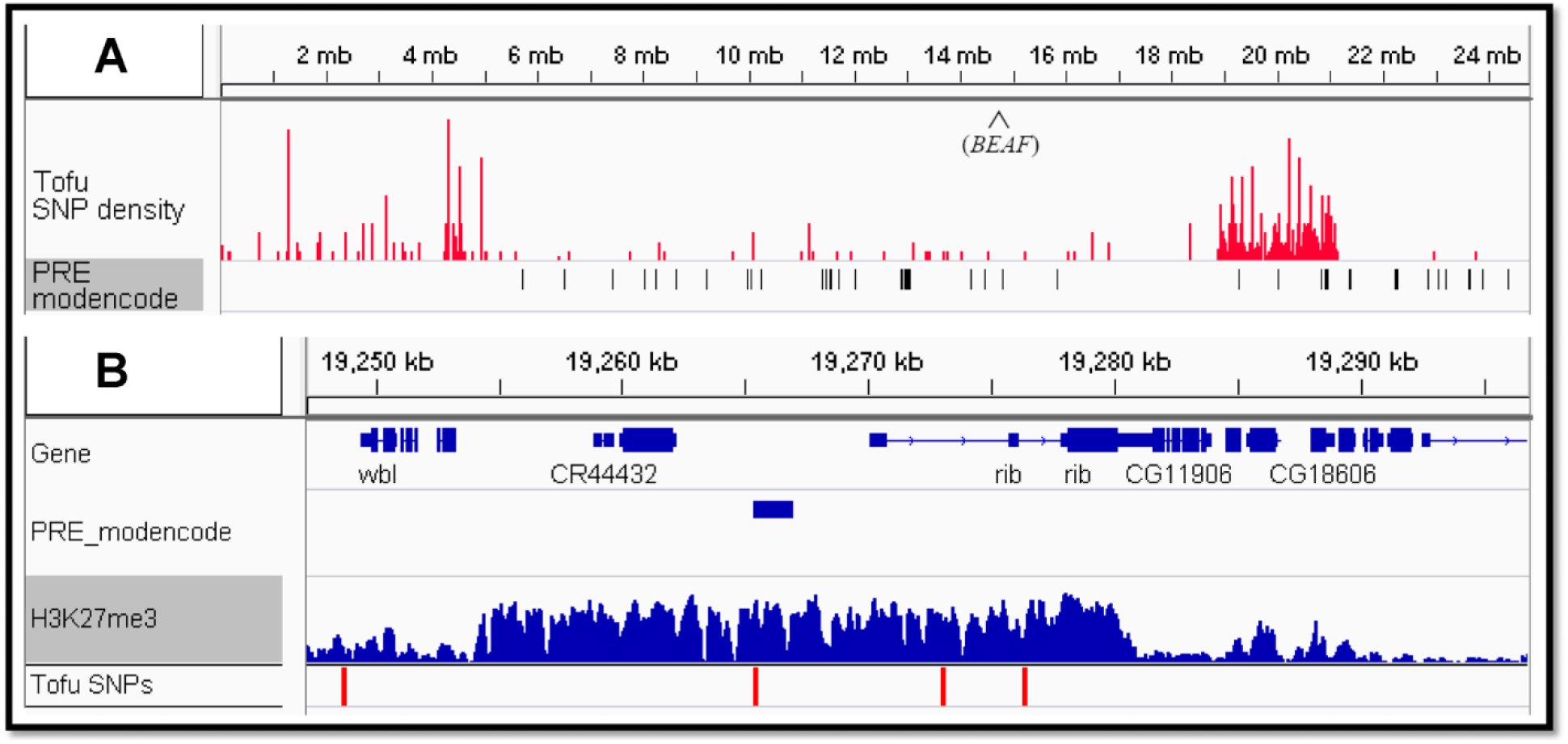
A SNP in a predicted PRE upstream of the *rib* gene is a candidate for the *Tofu* mutation. (A) IGV screenshot of chromosome 2R showing SNP distribution of *Tofu* candidates (red) and predicted PREs (black). Note the SNP island centered around 20 Mb. (B) IGV screenshot of the candidate PRE upstream of the *rib* gene. Data for the histone H3K27me3 mark associated with Polycomb repression is from ovaries (SRX177656). PRE SNP: 2R:19,265,447 (AATGCTGCTGCTCGA[T to G]CCTGACAAGCGTGTG)

Our lab previously performed a co-immunoprecipitation of BEAF from *Drosophila* embryos and by tandem mass spectrometry, identified around 100 co-immunoprecipitated factors with more than 1 peptide identified, a Benjamini-Hochberg corrected p-value less than 0.10, and a ratio of experimental to control IP greater than 1.0 (McKowen et al., 2022). One of these was *rib* (3 peptides, 1.8-fold enriched, 0.0390 p-value), making the SNP in the upstream PRE a candidate for *Tofu*. ChIP-seq data from ovaries (SRX177656) shows that the PRE upstream of *rib* is in a 25 kb island of the histone H3K27me3 mark that is associated with Polycomb silencing (Peng et al., 2016), suggesting *rib* is normally silenced in ovaries.

Consistent with this, we examined a single-cell atlas of gene expression in *Drosophila* ovaries (Jevitt et al., 2020) and found that *rib* is not expressed in any adult ovarian cell types. On the other hand, *rib* is involved in embryonic gonad development (Silva et al., 2016). It also has developmental roles in Malpighian tubules, salivary glands, hindgut, tracheae, and head (Jack and Myette, 1997). The PRE SNP could cause *rib* to be expressed in adult ovaries where it could compensate for loss of BEAF to rescue the fertility defect.

Additional considerations are consistent with the idea that misexpression of *rib* in ovaries could rescue the fertility defect caused by null *BEAF* mutations. ChIP-seq data from embryonic salivary glands identified 494 Rib-associated genes in the developing glands (Loganathan et al., 2016). We previously generated a list of 3241 BEAF-associated genes from S2 cell ChIP-seq data (Liang et al., 2014; McKowen et al., 2022), and found overlap with 30% (150) of the Rib-associated genes. A motif related to the BEAF-32B DNA binding consensus motif, CGATA (Hart et al., 1997; Zhao et al., 1995), represented around 15% of the occurrences of the top 10 motifs found in the Rib ChIP-seq peaks associated with the 494 genes (Loganathan et al., 2016). A more recent study found that Rib is associated with 73 of 84 ribosomal protein genes that are active in developing salivary glands and increases the expression of these genes, at least partially accounting for the requirement for Rib for salivary gland development (Loganathan et al., 2022). This work failed to detect sequence-specific DNA binding by Rib suggesting that it is recruited to DNA by other proteins. Our analysis of S2 cell BEAF ChIP-seq data found that BEAF is associated with 61 of these ribosomal protein genes, with both Rib and BEAF found at 53 of these genes. We also demonstrated that BEAF can activate a ribosomal protein promoter (*RpS12*) in a transient transfection assay (Dong et al., 2020). BEAF could be one of several factors that helps recruit Rib to promoters. Other regulators of ribosomal protein gene expression (TRF2, M1BP, DREF) can co-immunoprecipitate Rib, making them additional candidates (Loganathan et al., 2022). If the SNP in the predicted PRE directly upstream of the *rib* promoter leads to *rib* activation in adult ovaries, Rib could replace the function of BEAF at genes such as those encoding ribosomal proteins to largely restore ovary function.

## DISCUSSION

The fortuitous discovery of *Tofu* and finding that it is on chromosome 2, as is *BEAF*, led us to try to determine its location relative to *BEAF* by meiotic mapping. Our attempts at mapping were unsuccessful due to the difficulty of scoring fertility. Because the *BEAF*^*AB-KO*^ and *BEAF*^*NP6377*^ mutations were independently derived (Hayashi et al., 2002; Roy et al., 2007) but the *Tofu ABKO* and *Tofu NP* chromosomes were related by recombination, we decided to try using genomic sequencing to identify *Tofu*.

In retrospect, our genomic sequencing did not sample enough populations containing the *Tofu* mutation. Genome wide association studies in humans use many genomes from individuals to identify SNPs associated with a trait. These studies include a minimum of 2,000 individuals in each group, often exceeding 100,000 (Spencer et al., 2009; Tam et al., 2019). Our analysis was limited to 5 genomes of pooled populations, and this likely limited our ability to screen SNPs. However, similar pooled sequencing strategies have been employed successfully in *Drosophila* (Blumenstiel et al., 2009).

Because *Tofu* is a dominant mutation, we originally focused on SNPs that might cause a gain-of-function protein coding change. The lack of strong candidates led us to consider cis-regulatory element mutations. We also realized that SNPs near *Tofu* would likely remain associated with *Tofu*, so we looked for chromosome 2 SNP islands in our data. There was one island, and it contained 80% of the chromosome 2 SNPs. This did not narrow down the number of SNPs to consider very much, but it did localize a region that likely contains *Tofu*. CRE mutations and affected genes can be difficult to identify, but we considered a silencer element or repressive chromatin domain as likely candidates and took advantage of predicted PREs (Negre et al., 2011). Only 2 predicted PREs in the SNP island contained SNPs, leading us to focus on a predicted PRE just upstream of the *rib* gene. The region around this PRE, including *rib*, is in a chromatin domain enriched with the repressive histone H3K27me3 modification in ovaries (Peng et al., 2016), and *rib* expression was not detected in a single-cell atlas of ovary gene expression (Jevitt et al., 2020).

Different strategies can be used to determine whether the SNP in the predicted PRE allows *rib* expression in ovaries to rescue the fertility defect caused by null *BEAF* mutations. One approach would be to use Fluorescence In Situ Hybridization (FISH) to screen for atypical *rib* expression in *Tofu* ovaries. Another approach would be to use the GAL4 *UAS* system with different GAL4 drivers to determine if *UAS::rib* expression in different ovary cell types can rescue the *ABKO* mutation. Another possibility would be to use CRISPR-Cas9 to introduce or correct the PRE mutation and determine the effect on fertility in the presence of a null *BEAF* mutation. If it turns out that *Tofu* is not a *rib* ovary misexpression allele and the PRE SNP is not involved, the genome sequences we generated provide a valuable resource for identifying other candidates.

## Supporting information

VCF fild S1

VCF file S2

VCF file S3

VCF file S4

VCF file S5

VCF file S6

VCF file S7

## Acknowledgments

The authors thank Victor Corces for the *BEAF*^*NP6377*^*/CyO GFP* fly stock; Peter Klein for assistance with genome analysis; the Bloomington *Drosophila* Stock Center (bdsc.indiana.edu, NIH grant P40 OD-018537) for fly stocks; and FlyBase (flybase.org, NIH NHGRI grant 5U41HG000739-30) as an essential *Drosophila* resource. This work was supported by NSF grant 1244100 from the Division of Molecular and Cellular Biosciences (nsf.gov).

## Supplementary Files

fileS1_np63dm6.raw.snps.indels.vcf: VCF file for the original *BEAF*^*NP6377*^ fly stock with *Tofu* and at least one recessive lethal mutation.

fileS2_npnpdm6.raw.snps.indels.vcf: VCF file for the fly stock with *BEAF*^*NP6377*^ and *Tofu* but lacking any recessive lethal mutation.

fileS3_kokodm6.raw.snps.indels.vcf: VCF for the fly stock with the *BEAF*^*AB-KO*^ and *Tofu* alleles. fileS4_kog30dm6.raw.snps.indels.vcf: VCF for the original fly stock with *BEAF*^*AB-KO*^ but lacking *Tofu*.

fileS5_whitedm6.raw.snps.indels.vcf: VCF for the *y*^*-*^ *w*^*-*^ fly stock used as a wild-type genome. fileS6_tofu.vcf: VCF filtered to only retain variants unique to fly stocks with *Tofu* (present in *NP6377, Tofu NP*, and *Tofu KO* but not in *ABKO* or *y*^*-*^ *w*^*-*^).

fileS7_tofuchrom2.vcf: VCF with snpEff analysis of chr2 variants unique to fly stocks with *Tofu* (present in *NP6377, Tofu NP*, and *Tofu KO* but not in *ABKO* or *y*^*-*^ *w*^*-*^).

## Notes

https://www.ncbi.nlm.nih.gov/bioproject/?term=PRJNA1069327

### Competing Interest Statement

The authors have declared no competing interest.

